# T2T Gap-free Genome Assembly of Gp03, a Soybean Cultivar with High Genetic Transformation Efficiency

**DOI:** 10.1101/2025.07.21.665664

**Authors:** Wanjie Feng, Lijie Lian, Hongwei Xun, Dongquan Guo, Xutong Wang

**Author notes:** Corresponding author Email address (X. Wang); (D. Guo).

## Abstract

Soybean is a major cultivated crop worldwide, serving as a crucial source of edible oil, plant protein, and animal feed. The continuous growth of the global population necessitates accelerated soybean breeding. High-quality reference genomes are foundational for soybean functional genomics and breeding research. Several telomere-to-telomere (T2T) reference genomes have been released, facilitating subsequent functional studies. However, soybean varieties show varying efficiencies in genetic transformation, with currently available reference genomes having low transformation efficiencies. Here, we report the complete T2T genome assembly of Gp03, a soybean cultivar with high genetic transformation efficiency, developed in Northeast China. Gp03 exhibited high transformation efficiency in comparative studies and favorable agronomic traits for dense planting. The assembly comprises 20 chromosomes with a total size of 1.01 GB, an N50 length of 51.5 Mb, and includes 40 telomeres and 20 putative centromere regions. Genome annotation identified 61,832 genes and 545.8 Mb of repetitive sequences, accounting for 53.8% of the genome. This work provides a valuable genetic resource for soybean breeding and is expected to accelerate molecular breeding efforts.

---

Soybean (*Glycine max*) is a critical crop globally, contributing significantly to the production of edible oil, plant protein, and animal feed (Pagano and Miransari, 2016). With the global population projected to reach approximately 10 billion by 2050, there is an urgent need to accelerate breeding programs to ensure food security (Feeding the future global population, 2024). High-quality reference genomes are essential for advancing soybean functional genomics and breeding research. The first soybean reference genome for the cultivar Williams 82, released in 2010, has been a pivotal resource but suffers from low genetic transformation efficiency (Schmutz et al., 2010). Similarly, the ZH13 variety, widely cultivated in China, has been assembled and applied in research, yet also shows low transformation efficiency (Zhang et al., 2023). These limitations impede rapid functional genomic studies and breeding efforts.

To identify materials with superior genetic transformation efficiency, we conducted multiple tests on several soybean varieties frequently used in practical research. The results indicated that JishengP03 (Gp03), a soybean cultivar bred in Northeast China, exhibited significantly better transformation efficiency than other varieties (the parental lines) (Supplemental Table 1). This highlights Gp03’s potential in soybean functional genomics research and genetic improvement. Additionally, we examined the agronomic traits of Gp03. It displayed a small branching angle, making it an ideal plant type for dense planting (Figure 1A). Gp03 also has a shorter growth cycle compared to Wm82, which can reduce production cycles and save costs.

**Figure 1.**
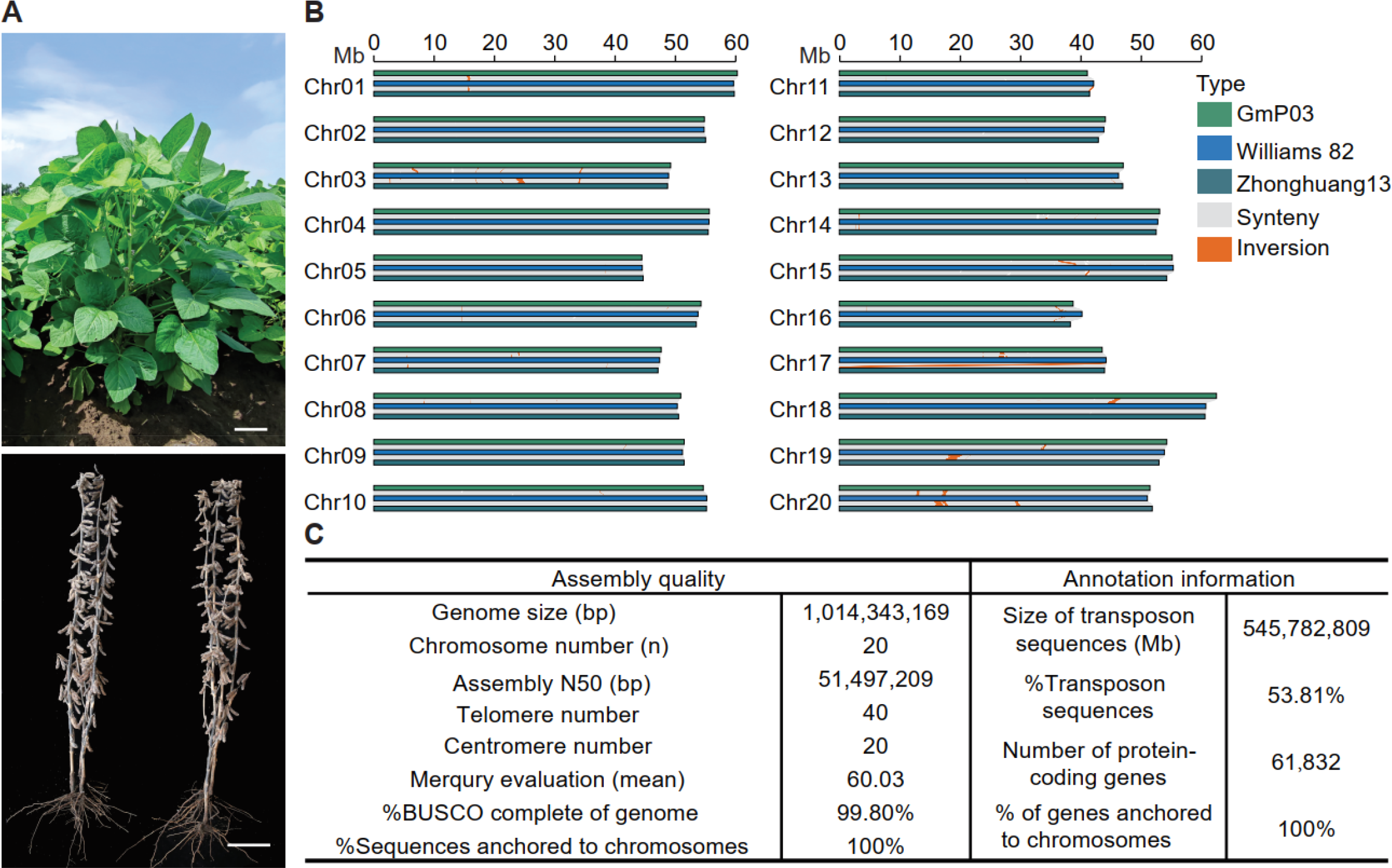
Field phenotype and genomic assembly summary of soybean material Gp03. (A) The appearance of Gp03 material at the immature stage (top) and the mature stage (bottom) in the field. (B) Genomic-wide alignment of Gp03 with Williams 82 and Zhonghuang13 (ZH13). (C) Statistics of genomic assembly indicators of Gp03

To effectively utilize Gp03 as a foundational genetic material, a well-assembled genome is essential. We employed multiple sequencing technologies to assemble a high-quality reference genome, generating high-coverage PacBio HiFi reads, Oxford Nanopore Technology (ONT) ultra-long reads, high-throughput chromatin conformation capture (Hi-C) sequencing data, and Illumina whole genome sequencing (WGS) data. These datasets provided the essential foundation for subsequent assembly. The N50 length of the HiFi reads was 18.5 kb, and the N50 length of the ONT reads was 100 kb (Supplemental Table 2). Hifiasm (0.19.8-r602) was used to integrate the HiFi and ONT reads, producing an initial assembly with a size of 1.01 Gb, an N50 length of 49.2 Mb, and comprising 73 contigs (Cheng et al., 2024).

To ensure the assembly’s accuracy, all small fragments were aligned to the NT database to remove sequences belonging to chloroplasts and mitochondria, which are not part of the nuclear genome. Following this, Hi-C reads were aligned to the genome using Juicer (version 2.2) software, and the results were processed to generate final alignments (Durand et al., 2016). These alignments were then fed into the 3D-DNA pipeline (201008) for chromosome-level scaffolding, producing 20 chromosome-level scaffolds (Dudchenko et al., 2017). Manual error correction was performed using Juicebox, and the results were reprocessed with 3D-DNA, resulting in a genome assembly with eight gap regions. Finally, we used the original HiFi and ONT reads for gap filling, and TGS-Gapcloser (version 1.2.1) successfully filled all gaps, resulting in a gap-free genome assembly (Xu et al., 2020) (Figure 1C).

Telomeres are repetitive DNA sequences at the ends of eukaryotic chromosomes, playing a crucial role in maintaining chromosome integrity and regulating the cell division cycle. Using tidk software, we identified telomeric repeat units in the genome, successfully discovering 40 telomeres. To locate the candidate centromere regions, we performed analysis using the online version of quartet (version 1.1.7) software (Lin et al., 2023). The analysis revealed that the average length of the centromeres across all chromosomes is 1,746,990 bp. The centromere regions averaged 1.75 Mb, with the longest on chromosome 20 (5,778,548 bp) and the shortest on chromosome 15 (598,647 bp). These findings provide insights into the genomic stability and chromosomal architecture of Gp03 (Figure 1C).

We conducted extensive validation of the assembled genome. The final genome consists of 20 chromosomes, with a total length of 1.01 Gb and an N50 length of 51.49 Mb. To determine the completeness of the genome assembly, we used the Benchmarking Universal Single-Copy Orthologs (BUSCO) score. The Gp03 assembly captured 99.8% of the Embryophyta BUSCO genes, indicating that 99.8% of the conserved core genes in plants are present in our genome (Figure 1C). Additionally, we aligned HiFi, ONT, and NGS reads to the genome, achieving alignment rates of 99.99%, 99.98%, and 99.58%, respectively. Furthermore, we used Merqury to assess the genome assembly accuracy with HiFi data, resulting in a score of 60.02. These results indicate that our Gp03 genome represents one of the highest quality assemblies reported for soybean to date, with a high degree of contiguity and base accuracy.

The Gp03 genome harbored 53.8% repetitive sequences, totaling 545,782,809 bp, predominantly long terminal repeat (LTR) retrotransposons, which accounted for 34.7% of the genome. Within the LTR category, Gypsy elements are the dominant type, accounting for 18.03%. In contrast, transposons like terminal inverted repeats (TIR) are relatively less abundant. To identify protein-coding genes, we integrated homology-based, de novo, and transcriptome-based prediction methods, identifying a total of 61,832 genes. The average length of protein-coding genes in the genome is 8,028 bp, spanning an average of 10 exons. Functional annotation of the genes was then performed using the eggNOG-Mapper online tool (Huerta-Cepas et al., 2019) (Figure 1C).

Whole-genome alignment revealed a high degree of collinearity between Gp03 and other soybean reference genomes, indicating the reliability of the assembly (Figure 1B). We also identified structural variations, including small-scale variations and large-scale structural variations such as duplications, inversions, and translocations. For example, we conducted a genome comparison with the Wm82-NJAU genome sequence and identified a total of 600 structural variants (SVs), which included 45 inversions, 138 translocations, and 220 duplications. Similarly, a comparison with the ZH13-T2T genome revealed 298 structural variants, including 61 inversions, 89 translocations, and 148 duplications (Figure 1B). Chromosomal inversions can hinder recombination in the inverted regions, impacting local adaptation and species differentiation. Inversions in genes related to regeneration ability may contribute to the distinct separation of Gp03 from other varieties, leading to its significantly higher receptor efficiency compared to other soybean cultivars.

Our study demonstrates that Gp03 is a valuable genetic resource for soybean breeding and functional genomics research. The high-quality T2T genome assembly of Gp03 offers a robust reference genome that can accelerate molecular breeding efforts and enhance our understanding of soybean genetics. The high transformation efficiency and detailed genomic information will facilitate targeted genetic modifications and breeding programs, ultimately contributing to the development of soybean varieties with improved yield, stress resistance, and other desirable traits. Moreover, the high-quality assembly of the Gp03 genome provides a foundation for exploring the functional roles of genes and regulatory elements in soybean. The superior genetic transformation efficiency and favorable agronomic traits of Gp03 make it an ideal model system for future studies aimed at improving soybean cultivation and addressing global food security challenges.

In conclusion, the T2T genome assembly of Gp03 represents a significant advancement in soybean genomics, offering a high-quality reference genome with superior genetic transformation efficiency. The detailed genomic information, including annotations of protein-coding and non-coding genes, structural variations, and chromosomal features, provides a comprehensive resource for soybean research. This work paves the way for advanced functional genomics studies and molecular breeding programs aimed at developing soybean varieties with improved agronomic traits and stress resistance. By leveraging the high transformation efficiency and favorable genomic features of Gp03, researchers can accelerate the development of soybean cultivars tailored to meet the demands of global food security. The genome, transcript, and protein sequences of Gp03 are available for BLAST search on our online server: http://xtlab.hzau.edu.cn/OnlineBlast/.

## Notes

### Competing Interest Statement

The authors have declared no competing interest.

